# Partitioning heritability using single-cell multi-omics identifies a novel macrophage subpopulation conveying genetic risks of coronary artery disease

**DOI:** 10.1101/2023.09.14.557845

**Authors:** Jiahao Jiang, Thomas K. Hiron, Thomas Agbaedeng, Yashaswat Malhotra, Edward Drydale, James Bancroft, Esther Ng, Michael E. Reschen, Lucy J. Davison, Chris A. O’Callaghan

**Affiliations:** Wellcome Centre for Human Genetics, Nuffield Department of Medicine, University of Oxford, Oxford, UK; Kennedy Institute of Rheumatology, Nuffield Department of Orthopaedics, Rheumatology and Musculoskeletal Sciences, University of Oxford, Oxford, UK; Oxford University Hospitals NHS Foundation Trust, John Radcliffe Hospital, Oxford, UK; Department of Clinical Science and Services, Royal Veterinary College, Hatfield, UK

## Abstract

**Background:** Coronary artery disease (CAD), the leading cause of death worldwide, is influenced by both environmental and genetic factors. While over 250 genetic risk loci have been identified through genome-wide association studies, the specific causal variants and their regulatory mechanisms are still largely unknown, particularly in disease-relevant cell types like macrophages.

**Methods:** We utilized single-cell RNA-seq (scRNA-seq) and single-cell multi-omics approaches in primary human monocyte-derived macrophages to explore the transcriptional regulatory network involved in a critical pathogenic event of coronary atherosclerosis—the formation of lipid-laden foam cells. Meta-analysis of scRNA-seq datasets from 26 human plaque samples was undertaken to provide a comprehensive atlas of lesional macrophages and to correlate subpopulations *in vivo* and *ex vivo*. The genetic risk levels of CAD were assessed by partitioning disease heritability across different macrophage subpopulations.

**Results:** We identified a novel macrophage subpopulation, termed lipid-handling macrophages, both *ex vivo* and *in vivo*, and identified associated marker genes, transcription regulators, and functional pathways. 18,782 cis-regulatory elements were identified by jointly profiling the gene expression and chromatin accessibility of >5000 macrophages. Integration with CAD GWAS data prioritized 121 CAD-related genetic variants and 56 candidate causal genes. We showed that CAD heritability was not uniformly distributed and was particularly enriched in the gene programs of lipid-handling macrophages. We investigated the cis-regulatory effect of a risk variant rs10488763 on *FDX1,* implicating the recruitment of AP-1 and C/EBP-beta in the causal mechanisms at this locus.

**Conclusions:** Our results provide genetic evidence of the divergent roles of macrophage subsets in atherogenesis and highlight lipid-handling macrophages as a key sub-population through which genetic variants actively influence disease. These findings provide an unbiased framework for functional fine-mapping of GWAS results using single-cell multi-omics and offer new insights into the genotype-environment interactions underlying atherosclerotic disease.

## Introduction

Coronary artery disease (CAD) is a leading global cause of morbidity and mortality [1] and results from atherosclerosis, a chronic inflammatory process [2]. Monocytes and macrophages play a central role in atherosclerosis and can engulf modified cholesterol-carrying lipoproteins to form lipid-laden "foam cells" within the lesions [3–5]. Cells of this lineage can display remarkably plastic responses to their microenvironment [6, 7], and human atherosclerotic lesions contain macrophages with pro-inflammatory “M1-like” to pro-resolution “M2-like” profiles [8–10]. Although single-cell techniques have demonstrated heterogeneity in human plaque macrophage populations [11–14], the lesions profiled are typically in the late stages of atherogenesis, when phenotypes may be polarized by long-established factors including modified cholesterols, cellular necrosis, sustained immune cell activation and concomitant cytokine release [4]. The response of macrophages to cholesterol, its vectors and modified derivatives is fundamental to atherosclerosis, but has not been well-defined at the single cell level and may differ across macrophage subpopulations [15, 16]. Deeper insight into the responses of different macrophage subpopulations to atherogenic lipid could help to identify new potential therapeutic targets.

CAD has a large heritable component [17], and more than 250 genomic risk loci have been identified by large genome-wide association studies (GWAS) [18, 19]. GWAS-identified variants are typically in linkage disequilibrium with multiple other variants and approximately 95% are in non-coding regions of the genome [20, 21]. However, despite considerable efforts [22], the cellular populations in which risk variants convey their disease risk and the molecular mechanisms whereby these loci transmit disease risk are largely unknown.

Advances in single-cell techniques provide new opportunities for investigating the function of non-coding variants by identifying the cis-regulatory interactions in cellular subpopulations [23–26], and using this information to prioritize candidate causal genetic variants [27–30]. However, transcriptome-based single cell approaches generally rely on public enhancer-gene maps. Since such maps are constructed mainly from bulk sequencing of immortalized cell lines or whole tissues, they do not fully capture gene regulation dynamics in in disease contexts. Variants prioritized based on single cell studies of chromatin accessibility alone have excess false positives, as only a subset of accessible chromatin sites have regulatory activity [31]. We reason that joint single-cell profiling of both the transcriptome and chromatin accessibility directly on primary cells will fill the gap between previous studies that only utilized one of the two modalities, providing a more comprehensive understanding of both the cellular and genetic components of human diseases.

In the current study, therefore, we undertook comprehensive single-cell RNA and multi-omics profiling of human macrophages to characterize their transcriptomic and epigenomic response to atherogenic oxidized low-density lipoprotein cholesterol (ox-LDL). We describe a novel human lipid-handling subpopulation in macrophages derived both from human atherosclerotic plaques and from circulating monocytes. Further, we constructed a high-resolution map of macrophage cis-regulatory elements, which allowed us to prioritize candidate causal CAD risk variants and to partition CAD heritability across macrophages subpopulations. This provides a foundation for functional dissection of the cellular contexts and the molecular mechanisms through which CAD risk variants act. As exemplification, we demonstrate how this approach provides mechanistic insight into the cis-regulatory effect of the CAD-risk SNP rs10488763 at the FDX1/RDX locus. Our results provide an unbiased framework for functional fine-mapping of CAD GWAS results through partitioning of heritability among macrophage subpopulations and deliver genetic evidence for the diverse and divergent influences of different macrophage subpopulations on atherosclerosis and its heritability.

## Methods

### scRNA-seq analysis of human plaque

Raw data of 29 human plaque scRNA-seq datasets reported in four studies were retrieved from Gene Expression Omnibus (GEO) using accession numbers GSE131778 [12] GSE155512 [32], GSE159677 [13], and GSE196943 [14].

Sequencing reads were aligned and counted using Cell Ranger (v6.0; 10x Genomics, Pleasanton, CA) with the GRCh38 reference genome. Low-quality cells were removed with a threshold of (detected genes <200 | detected reads < 500 | mt% > 20%). Potential doublets with more than 5000 genes detected were also removed. After batch effect correction with Harmony [33], cells were clustered in Seurat [34] with a resolution of 0.1. Three datasets that failed integration were removed from downstream analysis. Clusters were visualized using uniform manifold approximation and projection (UMAP) and cell types were annotated using SCSA with marker genes reported in [13, 35].

Based on the cell type annotation, myeloid cells were subset and re-integrated in Seurat. Datasets with fewer than 30 myeloid cells were excluded. In brief, raw counts were first normalized with SCTransform [36] accounting for the cell cycle scores, cells were then integrated using Seurat’s CCA integration workflow [34], and re-clustered at a resolution of 0.35 using the first 26 PCs. Macrophage sub-populations were annotated manually based on their marker genes and markers reported in the literature [37, 38]. Lipid-associated macrophages were re-clustered using first 19 PCs at the resolution of 0.35. Differential gene expression was tested using the Wilcoxon test. All statistical tests were repeated in down-sampled datasets to account for sample imbalance in tissue of origin (plaque vs adjacent normal tissue).

### Primary cell culture

Ethical approval for the study was obtained from the NHS Research Ethics Committee (South Central-Hampshire B, reference 13/SC/0392) and all participants provided informed consent.

CD14+ cells were isolated from peripheral blood of healthy adult volunteers using magnetic microbeads (Milltenyi Biotec, Bergisch Gladbach, Germany) according to the manufacturer’s protocol. *Ex vivo* macrophages were differentiated from CD14+ monocytes in R10 medium supplemented with 50 ng/ml human macrophage colony stimulating factor (M-CSF) (eBioscience, San Diego, CA) for 7 days. The medium was replaced at 3- and 5-days post-isolation.

### Ox-LDL preparation and cell treatment

LDL was purified from human plasma by isopycnic ultracentrifugation, then oxidized overnight using 25uM CuCl_2_, as described previously [39]. DPBS solution (Thermo Scientific, Waltham, MA) from the last round of dialysis was used as the control buffer. The concentration of ox-LDL was determined using the BCA protein assay (Thermo Scientific). On day 7 post-isolation, *ex vivo* macrophages were cultured for 48 hours in R10 media containing either 50 ug/mL ox-LDL or an equal volume of control buffer. For immunofluorescence studies, LDL was labelled overnight at 37⁰C with 300ug DiI per mg LDL, before purification by ultracentrifugation and oxidization using CuCl_2_.

### scRNA-seq analysis of *ex vivo* macrophages

Barcoded sequencing libraries of *ex vivo* macrophages and foam cells (n=4 donors) were generated from single cells using 10X Genomics Chromium system with 3’ chemistry (v2) according to the manufacturer’s protocol and sequenced on the Illumina HiSeq4000platform. Sequencing reads were aligned and counted using Cell Ranger as described above. After count normalization, single cell profiles were mapped to the plaque macrophage atlas using SCTransform and labelled using MapQuery in Seurat. Top 24 PCs were used in the initial clustering, and cells were clustered at a resolution of 0.35.

### Immunofluorescence staining

For fluorescence microscopy, CD14+ monocytes were differentiated at 100,000 cells/well in 8-well ibiTreat μ-slides (Ibidi, Gräfelfing, Germany) for 7 days, as described above, before treatment with DiI-labelled ox-LDL or control buffer. For detection of CHI3L1, cells were treated with protein transport inhibitor cocktail (eBioscience) overnight according to the manufacturer’s instructions. Treated macrophages were fixed at room temperature for 15 mins with 4% PFA in PBS, followed by permeabilization with 0.1% Triton X-100 (Thermo Scientific) and 0.2% Tween-20 (Roche) in PBS. Fixed cells were incubated in blocking buffer (1% BSA, 2% FCS, 0.3M glycine and 0.1% Tween-20 in PBS) at room temperature for 1 hour. Primary antibody staining was carried out at 4°C overnight with the following antibody dilutions in blocking buffer: CD52-APC (clone HI186, BioLegend) 1:200, CHI3L1 (MM0628-9X24, Abcam) 1:500, MITF (HPA003259, Atlas Antibodies) 1:100 and tubulin-AF488 (clone DM1A, eBioscience) 1:500. For detection of the CHI3L1 and MITF primary antibodies, cells were incubated with FITC anti-rat IgG (Sigma-Aldrich) and AF633 anti-rabbit IgG (Invitrogen), respectively, at 1:1000 in blocking buffer in the dark at room temperature for 1 hour. Cells were washed three times with PBS-0.1% Tween-20 for 5 mins between stains. Nuclei were counterstained with Hoechst 33342 (Santa Cruz Biotechnology) diluted 1:10,000 in PBS. Stained cells were imaged on a LSM 900 with Airyscan2 in confocal mode (Zeiss). Quantitative analysis was performed using arivis Vision4D version 4.1 (Zeiss).

### Single cell multi-omics

Sequencing libraries from *ex vivo* macrophages and foam cells (n=4 donors) were generated from single nuclei using 10X Genomics Multiome system according to the manufacturer’s instructions and sequenced on the Illumina NovaSeq6000 platform. Reads were aligned and counted using Cell Ranger ARC (v2; 10x Genomics) with the GRCh38 reference genome. For snATAC-seq, peaks were trimmed and re-called with MASC3 [40]. Count matrices were processed with Seurat (v4.0.4) and Signac (v1.4.0). Low-quality cells and potential doublets were filtered out with the threshold of (TSSE <1 | detected gene/peak < 1000 | detected gene/peak > 50,000).

For RNA analysis, counts were normalized with SCTransform and integrated using Seurat’s CCA integration workflow. For ATAC analysis, count matrices were normalized in Signac with default parameters and batch corrected using Harmony. Cells were clustered using the Louvain algorithm at a resolution of 0.3 and were projected using UMAP based on the top 26 RNA reduced dimensions and top 10 ATAC reduced dimensions, following Seruat’s WNN workflow [41].

Cluster marker genes were identified by a logistic regression test with mitochondrial percentage as a latent variable, and p-values were adjusted using the Benjamini-Hochberg procedure. KEGG and GO-BP enrichment was performed using XGR [42]. Enriched GO terms were reduced and visualized using REVIGO [43]. Differential abundance of sub-populations between ox-LDL and buffer samples was analyzed using a t-test.

### TF motif deviation analysis

After generating background peak sets matching GC content and average accessibility, motif deviation scores were computed using ChromVAR [23]. Human TF motifs were downloaded and manually curated from cisBP, as described in [44].

### Cis-regulatory analysis

Peak to gene correlation was calculated as described [41, 44]. The peak-gene correlation coefficient was calculated using the mean-centered ATAC-seq signals for a given peak and a corresponding gene expression. Statistical significance of a given peak-gene pair was estimated from a null distribution computed using a set of background peaks matched for GC content and average accessibility. CREs were annotated using the ChIPseeker R package [45].

For DORC analysis, genes with at least 9 identified CREs were selected as DORCs. By summarizing the number of reads mapped to the CREs of each DORC gene, the snATAC-seq peak matrix was transformed into a DORC score matrix. The DORC-TF regulatory network was constructed as described [44]. In brief, DORC genes were pooled with their k-nearest neighbors based on the DORC scores (k=30). The enrichment of each TF motif was computed using the CREs of each DORC gene pool over the matched background peaks. The smoothed RNA expression of each TF tested was correlated with DORC-scores in all cells, and the significance tested using standardized Spearman correlation levels. A regulatory score per DORC-TF pairs was calculated by combining two significance levels (motif enrichment and correlation). Network plots for CAD-DORCs (DORC genes with CREs overlapping CAD-related SNPs) were drawn for the strongest associations (score > 1.5) using the ggnet2 R package.

### ChIP-seq data analysis

H3K27ac and CEBPB ChIP-seq datasets were generated previously and retrieved from GEO (GSE54975) [39]. For H3K27ac, reads from two biological replicates were aligned to GRCh38 using Bowtie2 [46] and merged into a single bam file per condition. PCR duplicates were marked and removed using GATK MarkDuplicates. Signals were normalized and plotted using deepTools (v3.3.1). Smoothed conditional means were computed to evaluate the enhancer signals over CRE peaks using a generalized additive model in ggplot2.

### QTL SNPs

eQTL SNPs were obtained from GTEx v8 (whole blood) and eQTLGen (Blood) database. SNPs at hg19 coordinates were converted to hg38 using annotations from NCBI dbSNP Build 151. Chromatin accessibility QTL SNPs (human iPSC-derived macrophage) were downloaded from supplementary materials at the publisher’s website [47].

### GWAS – related SNPs

GWAS statistics were curated by NHGRI-EBI Catalog and downloaded from the UCSC hg38 database (April 2022). We identified SNPs with significant genome-wide association (p<5×10^−8^), pruned the list (r^2^ > 0.1 identified by 1000 Genome Project phase 3 in EUR samples, hg38 build) to remove redundant loci, and then pooled all SNPs in high LD within each locus (r^2^ > 0.8 in EUR samples) to establish a set of trait-associated variants. Only dbSNP common variants (MAF>1%, dbSNP155) were considered in this study. LD information was extracted using PLINK v1.90b6.26 [48].

### Enrichment analysis

We used chromVAR to create background peaks matched for GC content and chromatin accessibility levels across cells for each open chromatin region. For each annotation considered, SNPs/annotated regions were intersected with peaks using the findOverlaps R function, and the null distributions were constructed with n=1000 background peak sets. The significance of each trait-region overlap was determined using Z-statistics computed from the background null distributions.

### Stratification of CREs

We stratified cis-regulatory elements for each macrophage subpopulation by applying a logistic regression test for differential expression between subpopulations (one vs the rest), between treatment (ox-LDL vs buffer) of all cells, or between treatment within a subpopulation. The p values for genes in each test were then transformed to the relative weights (gene scores) as described previously [49]. In brief, gene scores were computed by min-max normalizing X scores, which follow a X^2^ distribution and were calculated as X = −2 log (p). Each CRE was then weighted by the sum of the scores of their target genes.

### LD score regression

LDSC v1.01 was used to partition CAD heritability and test the enrichment for each subpopulation as described [50, 51]. In brief, all 1000G SNPs were annotated and weighted by their overlaps with subpopulation CREs or subpopulation-ox-LDL CREs; we then inferred the LD score for each SNP using the 1000G EUR reference panel, and computed the heritability associated with each subpopulation using a multivariate linear regression model, conditioned on background annotations; the standard errors of enrichment statistics and regression coefficients are computed using a block jack-knife (n=1,000) [50]. The full summary statistics for CAD used in this study were downloaded from the GWAS Catalog, study ID GCST005194 [18].

We evaluated the extent and the significance of heritability enrichment using two metrics: the relative enrichment score *E_c_* over background annotations and the standardized effect size *τ*\*, as implemented by the LDSC software [51].

*E_c_* follows the normal distribution and is defined as: 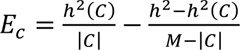, where *h*^2^(*C*) and *h*^2^ are the heritability of SNPs in category *C* and all SNPs respectively, *M* is the total number of SNPs included, and |*C*| is the number of SNPs in *C*. The relative enrichment over background was calculated by comparing *E_c_* to the background enrichment score *E_background_* constructed using either all macrophage open chromatin regions, the pseudo-bulk oxLDL-response program, or the pseudo-bulk disease-vs-healthy program.

For *ex vivo* macrophage and plaque macrophage subpopulation programs, we computed *τ*\* conditioned on all other subpopulations but the subpopulation of interest using a multivariate linear regression model. For subpopulation-ox-LDL/disease programs, we first constructed a pseudo-bulk ox-LDL/disease program, then computed the *τ*\* conditioned on this pseudo-bulk program for each subpopulation.

### Allelic analysis

Single-cell ATAC-seq libraries were aggregated by sample and genotyped using the GATK (v4.1.7.0) germline short variant discovery pipeline [52]. PCR duplicates were marked with MarkDuplicates. base quality scores were recalibrated against dbSNP155 known sites, and variants were called with HaplotypeCaller, combined using GenomicsDBImport, and genotyped with GenotypeGVCFs. Genotyping results from SNPs in perfect LD (r^2^=1 in 1000G EUR samples) were merged. Plaque scATAC-seq reads were obtained from GEO using accession number GSE175621.

To test the allele-specific enhancer activities in response to ox-LDL, *ex vivo* macrophages were transfected with pGL4.27 Firefly luciferase reporter plasmid and pRL-SV40 Renilla luciferase plasmid (1/50 DNA amount compared to Firefly, to control for transfection efficiency) using the Human Macrophage Nucleofector kit (Amaxa, Lonza, Portsmouth, NH). After 24 hours cells were treated with 50 ug/ml ox-LDL or buffer for a further 24 hours. Cells were lyzed and assayed for luciferase activity using the Dual Luciferase Reporter Assay System (Promega, Fitchburg, WI). Firefly luciferase activity was normalized to Renilla luciferase activity as relative light units (RLU). Two dsDNA constructs were synthesized (IDT gBlocks™) for each allele, one covering −75bp to +75bp, and one covering −100bp to +700bp of the targeted variant (FigS8E and Table S6). All inserts were verified by Sanger sequencing prior to transfection.

The coding sequence of the LAP isoform of CEBPB was cloned into the pcDNA3.1 vector, and transfected into *ex vivo* macrophages using the Nucleofector kit. After 24 hours cells were treated with 50 ug/ml ox-LDL or buffer for a further 24 hours, then lysed for RNA extraction.

The sequences of dsDNA constructs and qPCR primers for *FDX1*, *CEBPB*, and *GAPDH* were provided in Supplemental Notes.

## Results

### Monocyte-derived macrophages resemble the transcriptomic heterogeneity of the lipid-associated macrophage in atherosclerotic plaques

We created an atlas of macrophage subpopulations in human atherosclerotic plaques by meta-analysis of single-cell RNA-seq datasets from 26 human coronary and carotid artery plaques [12–14]. We first mapped all plaque cells (n= 86,275) to 8 broad cell types (FigS1A and S1B), and then focused on myeloid cells (n= 12,538), which were identified based on the expression of *CD14*, *CD68* and *AIF1*. Re-clustering of the myeloid cells identified heterogeneous myeloid populations in atherosclerotic plaques (Fig1A, FigS1C) including cycling macrophages, CD1+ dendritic cells, IGSF21+ macrophages, and two pro-inflammatory clusters (S100A8^hi^ macrophages, IFNG-activated macrophages) [35, 38]. We also identified a population of ‘SEM’ (stem cell, endothelial cell, monocyte)-like cells, with high expression of marker genes characteristic of a recently identified vascular smooth muscle cell-derived intermediate population [32]. Notably, two macrophage populations expressed lipid-associated markers (*SPP1*, *CD9*, *FABP5*): a TREM2+ lipid-associated macrophage (LAM) population similar to that identified in adipose tissue [37], and a TREM2-/CCL7+ macrophage population with high pro-inflammatory potential [53].

**Fig1.**
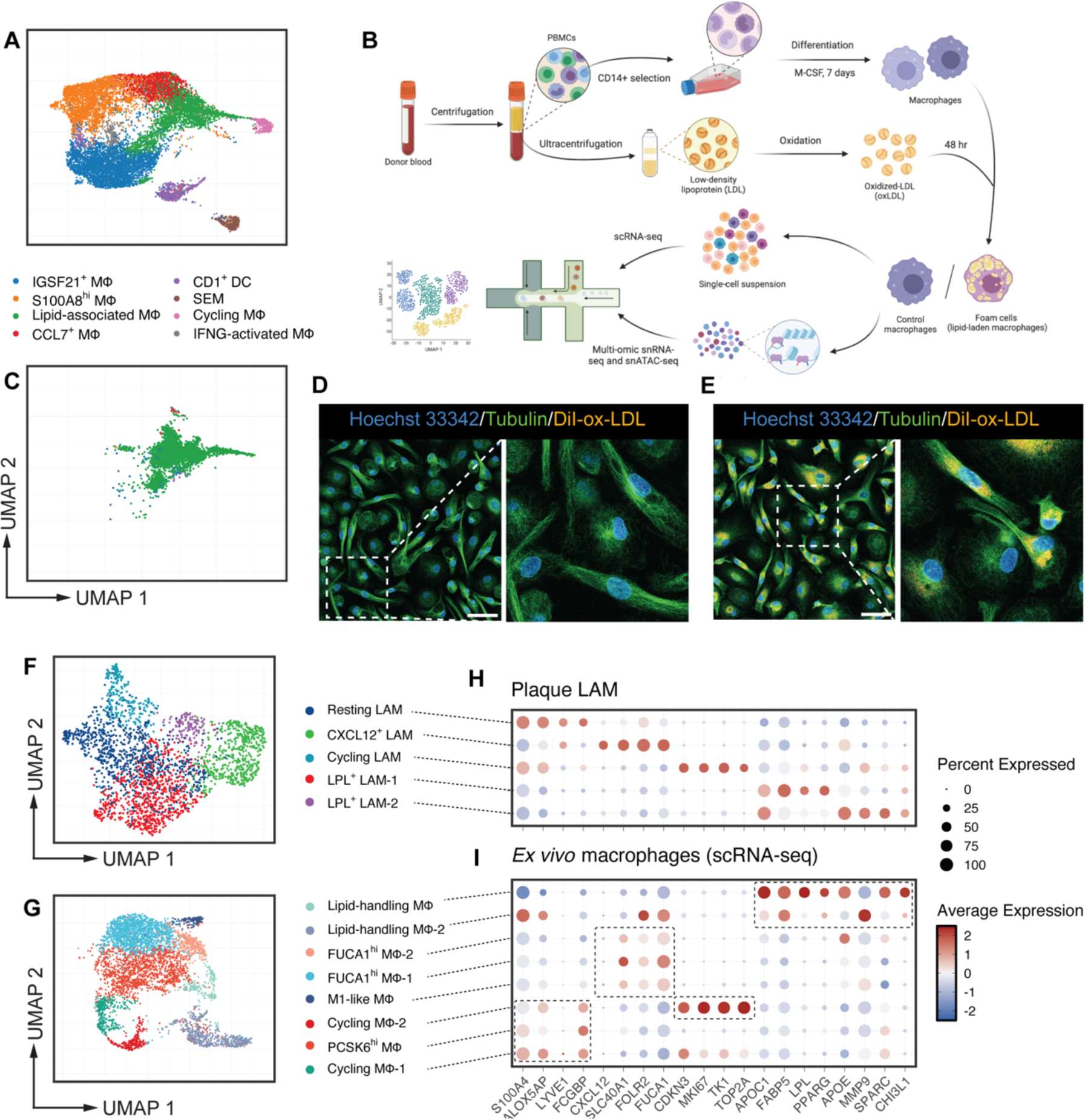
Human *ex vivo* macrophages map to lipid-associated macrophages in plaque. **A**, The plaque macrophage atlas: multiple subpopulations of plaque macrophages demonstrated by meta-analysis of plaque scRNA-seq datasets [12–14], and displayed using uniform manifold approximation and projection (UMAP). **B**, Experimental strategy for single-cell transcriptomic and multi-omics analyses. **C**, Single-cell transcriptomic profiles of *ex vivo* macrophages projected onto the plaque macrophage atlas (**A**) and displayed using UMAP, demonstrating close mapping to the lipid-associated macrophages subpopulation in plaques. Color represents the predicted population as in (**A**). **D** and **E**, Representative immunostaining of control (**D**) and lipid-laden *ex vivo* macrophages (**E**), showing variations in cellular morphology and lipid accumulation. Scale bar: 50μm. **F** and **G**, UMAP visualization of re-clustered subpopulations of the lipid-associated macrophage (LAM) (**F**) and the *ex vivo* macrophage (**G**) populations demonstrating macrophage heterogeneity at the sub-population level. **H**, Gene expression patterns of selected marker genes for different plaque LAM subpopulations. Color denotes scaled average expression and size denotes the percentage of the population expressing the gene. **I**, Gene expression patterns of the markers in (**H)** across *ex vivo* macrophage subpopulations*. Ex vivo* subpopulations displaying similar expression profiles are grouped in dotted box. Markers of plaque LPL^+^ LAM subpopulations are also highly expressed in lipid-handling macrophages.

Monocytes adhere to the vessel wall at sites of atherosclerotic lesions and have been widely recognized as the predominant source of plaque macrophages [3, 4, 54]. Primary macrophages derived *ex vivo* from circulating monocytes are extensively used to study the molecular basis of atherosclerosis, especially the uptake of atherogenic lipid, such as oxidized low density lipoprotein cholesterol and the consequent development of lipid-laden macrophages (foam cells) [39, 55-57]. However, how these macrophages map to *in vivo* plaque macrophages is not clearly defined. To address this, we profiled the transcriptomic changes induced by atherogenic ox-LDL in primary human monocyte-derived macrophages at single-cell resolution (Fig1B). For clarity, we term myeloid cells derived from human plaques as plaque macrophages and macrophages derived from circulating monocytes as *ex vivo* macrophages, respectively. Single-cell RNA-seq profiles of *ex vivo* macrophages (n=4,841) were integrated and reference-mapped onto our atlas of plaque macrophages. Both unexposed and ox-LDL-exposed *ex vivo* macrophages mapped tightly to the TREM2+ LAM presented in the atherosclerotic plaques (Fig1C and FigS1D). This cellular identity was further evidenced by high expression of LAM marker genes [37, 58] across all *ex vivo* macrophages (FigS1E).

Fluorescence microscopy demonstrated extensive heterogeneity in cell morphology, lipid-accumulation and lipid distribution among *ex vivo* macrophages (Fig1D and 1E). As these macrophages mapped to the LAM population in plaques, we hypothesized that plaque LAMs would also be heterogeneous with functionally distinct subpopulations. We therefore re-clustered plaque LAMs (n=2,487) and *ex vivo* macrophages separately using an unsupervised clustering approach at the same resolution to avoid over-clustering (Fig1F and 1G). This led to the identification of multiple distinct subpopulations in plaque LAM, including cycling cells (*CDKN3*, *MKI67*), resting LAM (*S100A4*, *ALOX5AP*), CXCL12+ LAM (*CXCL12*, *FUCA1*), and two LPL+ clusters (*LPL*, *CHI3L1*).

The *ex vivo* macrophage subpopulations had similar transcriptomic profiles to these plaque LAM subpopulations, although several markers (e.g., *CXCL12*) were expressed at higher levels in the plaque cell (Fig1H and 1I). Two *ex vivo* subpopulations showed particularly strong and cluster-specific expression for markers genes of LPL+ LAMs (Fig1I). We term these *ex vivo* macrophage subpopulations ‘lipid-handling macrophages’ because many of their marker genes are involved in lipid processing and metabolism. The gene expression patterns shared between the *ex vivo* lipid-handling macrophages and plaque LPL+ LAMs were not observed in other plaque macrophage populations (FigS1C), suggesting that activation of lipid metabolism might be a feature specific to LAMs. This further supports the link between *ex vivo* macrophages and plaque LAMs.

### Paired snRNA-seq and snATAC-seq recovers cellular heterogeneity in gene expression and transcription factor activation

To further characterize the cellular identities of macrophage subpopulations, we undertook single-cell multi-omics, simultaneously characterizing the transcriptome and open chromatin profile of each cell, using single-nucleus RNA-seq and ATAC-seq respectively (Fig1B). The single-nucleus RNA-seq profiles were highly concordant with the single-cell RNA-seq profiles (FigS2A). By integrating the single-nucleus RNA-seq and ATAC-seq data for each cell (n=5,478), we improved the resolution for the allocation of macrophage subpopulations (Fig2A-B, FigS2B, TableS1), and with this improved analysis, lipid-handling macrophages identified in the scRNA-seq data were clearly characterized as a single subpopulation (*CHI3L1*, *APOC1* and *LPL*). Other subpopulations included an M1-like population (*NFKBIA*, *ICAM1* and *CCL2*), two proliferating populations (*TOP2A*, *CENPA* and *CENPE*), and a small AK5+ population (*ANKH*, *AK5* and *MYO1B*) with some similarity in gene expression to osteoclasts [59]. Subpopulations analogous to classic resting M0 macrophages, included a PCSK6^hi^ population, (*PCSK6*, *ITGB5* and *SHC3*), two FUCA1hi populations (*FUCA1*, *WWP1* and *SELENOP*), and a transitional monocyte/macrophage FCGR^hi^ population (*FCGR3A*, *FCGR2A* and *PCSK6*) (Fig2A-B).

**Fig2.**
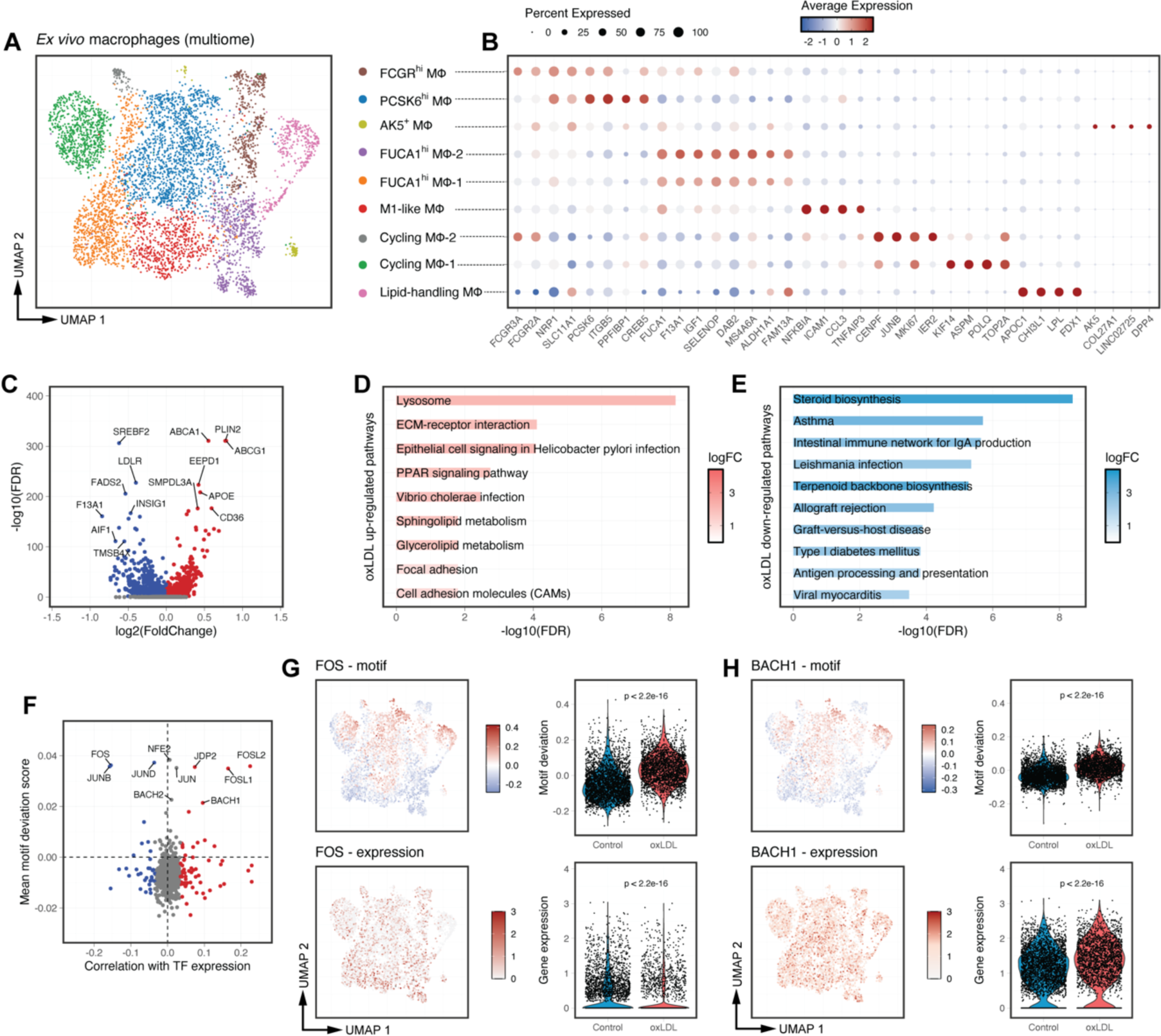
Single-cell multi-omics profiling of gene expression and chromatin accessibility in human macrophages. **A**, *Ex vivo* macrophage subpopulations based on integrated RNA-seq and ATAC-seq data from single cells and displayed as UMAP. **B**, Scaled average expression of marker genes for subpopulations in (**A**). **C**, Volcano plot showing the gene expression changes induced by ox-LDL. Genes with FDR < 0.1 are plotted with their log fold-changes (red for upregulation and blue for downregulation). **D** and **E**, KEGG pathways upregulated (**D**) or downregulated (**E**) in the ox-LDL-treated cells compared to buffer-treated cells. **F**, Mean TF motif deviation scores (y axis, the relative enrichment compared to background) plotted against their correlation with TF RNA expression (x axis, Pearson coefficient) in ox-LDL-treated cells. TFs are colored by significance (FDR < 0.1) for negative (blue) or positive (red) correlation. **G**, Motif deviation scores (top) and RNA expression (bottom) across for AP-1 family TF *FOS*. **H**, Motif deviation scores (top) and RNA expression (bottom) for NFE2 family TF *BACH1*. P-values were calculated using a Wilcoxon test.

To explore the transcription factors (TFs) activated in the different macrophage subpopulations, we assessed TF binding activity in individual cells by computing motif deviation scores, which represent motif-specific gain or loss of chromatin accessibility relative to the average cell profile (see Methods) (FigS2C). Since TFs belonging to the same family usually share binding motifs, we also examined the correlation between the motif deviation scores and TF expression to understand the role of TFs within the same family. As expected, TFs from the NF-kB family displayed both higher gene expression and higher motif deviation scores in M1-like macrophages (FigS2D-F).

### Single-cell multi-omics refines the transcriptomic and epigenomic response to ox-LDL in macrophages

Across all *ex vivo* macrophages, exposure to ox-LDL significantly upregulated the expression of genes involved in lipid transport including *CD36*, *ABCA1* and *ABCG1* (Fig2C, TableS2)[60, 61]. Kyoto Encyclopedia of Genes and Genomes (KEGG) and Gene Ontology Biological Process (GOBP) pathway analysis demonstrated that ox-LDL induced lysosomal activation and inhibited cholesterol biosynthesis (Fig2D-E, FigS2G-H).

Using bulk approaches, we have previously identified enrichment of the binding motifs of AP-1 and NFE-2 factors in dynamic enhancer sites following exposure of human macrophages to ox-LDL [39]. Both AP-1 factors and NFE-2 factors respond to oxidative stress [62, 63]. However, it is challenging to pinpoint the specific TFs involved in the oxLDL-response in macrophages, as they all share similar binding motifs. Here, with single cell multi-omics, we can now dissect the directional influence of individual AP-1 and NFE-2 family member TFs. For example, expression of two AP-1 family members *FOSL1* and *FOSL2* positively correlated with the chromatin accessibility of AP-1 motifs, while another AP-1 factor *FOS* displayed a reverse correlation (Fig2G). Expression of the NFE-2 family member *BACH1* was positively correlated with its motif binding (Fig2H), but no significant correlation was observed for other NFE-2 family members.

### Lipid-handling macrophages display activation of genes and transcription regulators involved in fatty acid metabolism

Lipid-handling macrophages, although comprising only 6.5% of the total *ex vivo* macrophage population (FigS3A), contributed significantly to the overall expression of ox-LDL-induced genes (e.g., 44.8% of total *APOC1* expression, Fig3A, TableS2). Therefore, we sought to further characterize these cells.

**Fig3.**
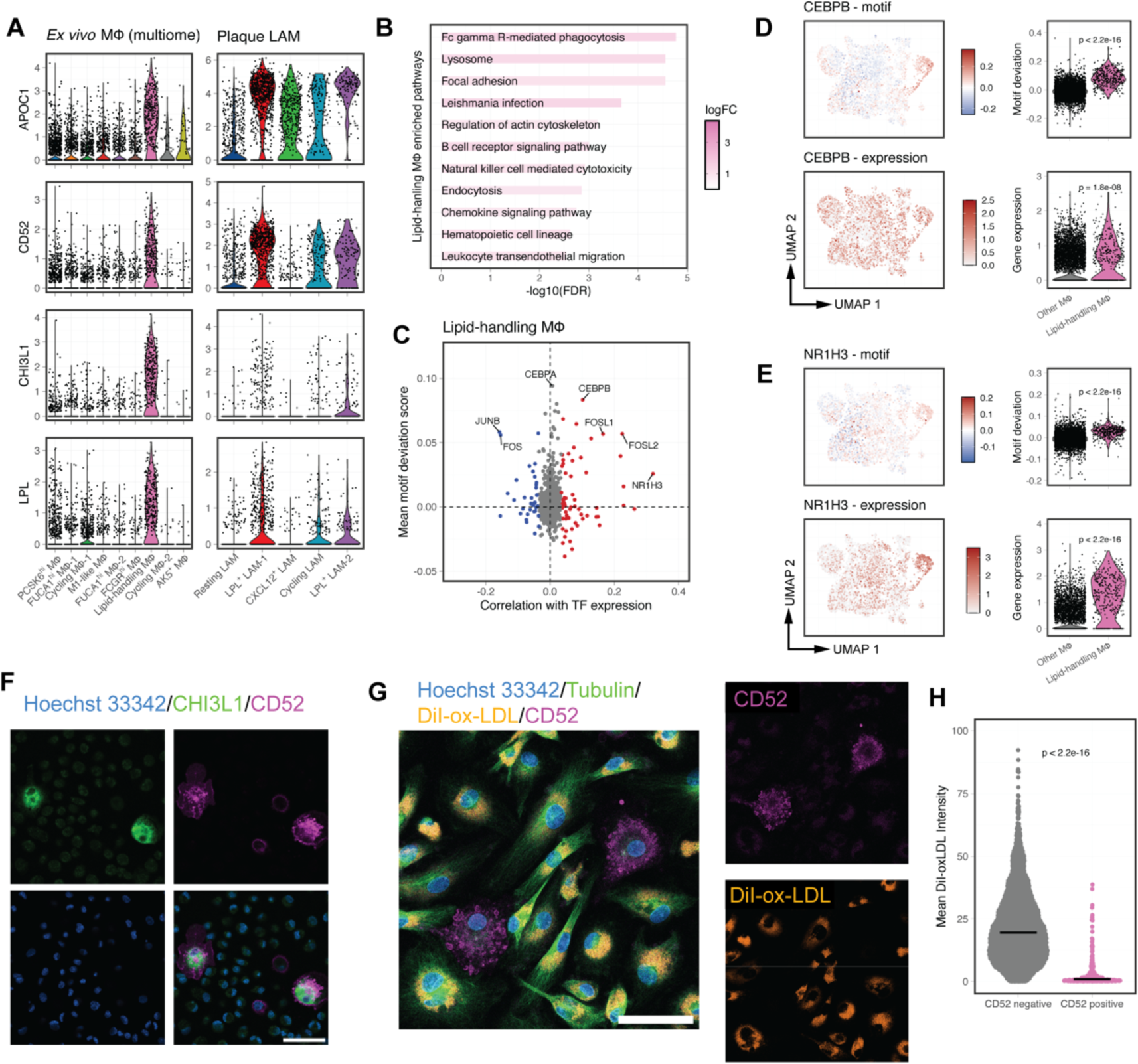
Multi-omics characterization of the lipid-handling macrophage subpopulation. **A**, Comparison of the expression of lipid-handling markers in *ex vivo* macrophages and plaque LAM subpopulations. **B**, KEGG pathways enriched in the marker genes of lipid-handling macrophages. **C**, Mean TF motif deviation scores (y axis, the relative enrichment compared to background) compared with their correlation with TF RNA expression (x axis, Pearson coefficient) in lipid-handling macrophages. TFs are colored by significance (FDR< 0.1) for negative (blue) or positive (red) correlation. **D**, Motif deviation scores (top) and RNA expression (bottom) for *CEBPB*. **E**, Motif deviation scores (top) and RNA expression (bottom) for *NR1H3*. **F**, Representative immunostaining of *ex vivo* macrophages for lipid-handling macrophage markers CHI3L1 and CD52. Scale bar: 50μm. **G**, Representative immunostaining of *ex vivo* macrophages for lipid accumulation after 48 hr treatment with DiI-ox-LDL. Scale bar: 50μm. **H**, Quantification of the intracellular level of DiI-ox-LDL in (**G**), shown as individual data points along with the median (arbitrary units). n=3091 cells from 3 biological donors studied. P-values were calculated using a Wilcoxon test.

*APOC1*, *CD52*, *CHI3L1* and *LPL* were among the most significant marker genes for lipid-handling macrophages in both our scRNA-seq and multi-omics data, and were also highly expressed in plaque LPL+ LAMs (Fig1I and Fig3A). Both *APOC1* and *LPL* are involved in lipoprotein metabolism [64]. Additionally, KEGG and GOBP enrichment analyses suggested that these macrophages have enhanced phagocytosis (Fig3B) and fatty acid metabolism (FigS3B). Notably, the marker genes of lipid-handling macrophage (133 genes with FDR < 0.1 and log2FC > 1, TableS1) were significantly overrepresented in a subcutaneous-fat and liver gene regulatory network (GRN51, FDR=1.42×10^−20^, FigS3C). The expression of this network strongly correlates with BMI and blood triglyceride levels (FigS3D) in a CAD patient cohort [65].

We then determined the transcriptional regulators specific to lipid-handling macrophages. CEBP family TFs showed the largest motif deviation score that was specific for lipid-handling macrophages (Fig3C). The expression of *CEBPB* was higher in lipid-handling macrophages and correlated positively with CEBP motif binding activity (Fig3D). There was also a strong correlation between expression and motif usage for the transcription factor NR1H3, a key regulator of genes involved in fatty acid metabolism (Fig3C and Fig3E) [66].

To functionally validate these findings, we first used immunofluorescence to confirm key marker gene expression and showed that CD52 was expressed exclusively in CHI3L1 positive cells (Fig3F). Next, we performed a lipid-uptake assay using DiI-labeled oxLDL which demonstrated reduced accumulation of intracellular ox-LDL in CD52+ macrophages compared to CD52-macrophages (Fig3G and Fig3H). After oxLDL treatment, lipid-handling macrophages exhibited increased expression of *CD36* (FigS3E), a well-documented macrophage receptor for oxLDL uptake [67]. The same upregulation of *CD36* was observed in other subpopulations and in bulk macrophages after oxLDL exposure (FigS3E and FigS3F), indicating that lipid-handling macrophages recognize and uptake oxLDL similarly to other subpopulations.

In summary, lipid-handling macrophages exhibit enhanced gene expression and TF activity related to lipoprotein metabolism. They have reduced intracellular lipid accumulation upon oxLDL exposure compared to other macrophage subpopulations, suggesting efficient lipid/cholesterol processing and clearance capacities.

### Analysis of cis-regulatory elements identifies complex CAD-associated transcriptional regulatory networks

Single cell multi-omics data allowed us to identify cis-regulatory interactions by correlating chromatin accessibility at non-coding peaks with expression of nearby genes (<500 kb) [26]. We identified 18,782 unique cis-regulatory elements (CREs) spanning 6,381 genes (TableS3). Of these CREs, 2,642 were correlated with expression of more than one gene, resulting in a total of 22,555 peak-gene pairs. The major locations of CREs were in introns (42.3%), promoters (24.4%), and intergenic regions (24.3%), with 42.6% of all CREs located >10kb from any recognized transcription start site (Fig4A).

**Fig4.**
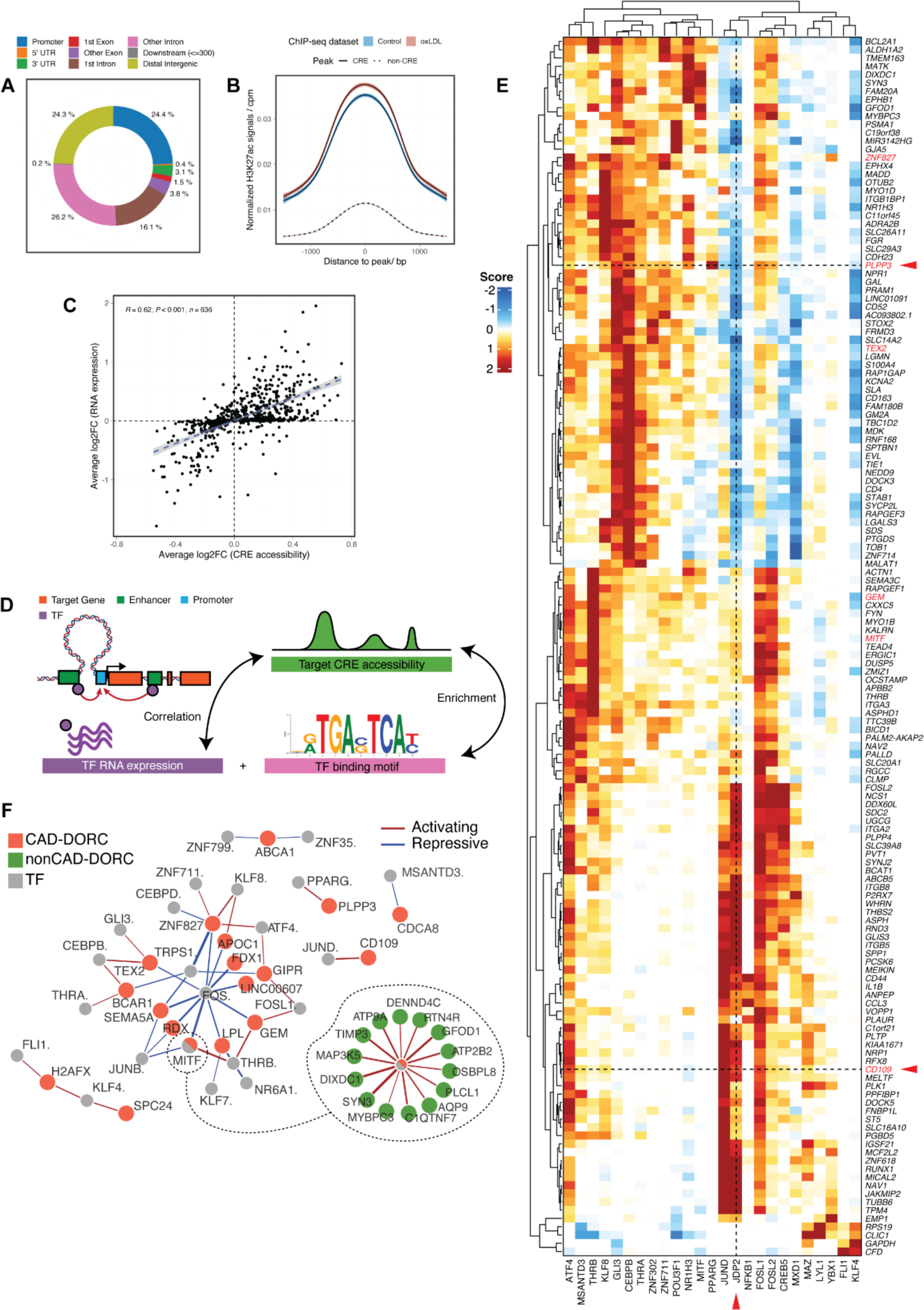
Multi-omics analysis identifies cis-regulatory elements and key transcriptional regulatory networks for CAD. **A**, Location of cis-regulatory elements (CREs) (n= 22,555 CREs). **B**, Aggregated H3K27ac signals at CRE-peak and non-CRE-peak regions. Signals from untreated (blue) and oxLDL-treated (red) human macrophages are plotted separately. Shaded regions indicate 95% confidence intervals. **C**, Correlation between ox-LDL-induced changes in CRE chromatin accessibility and in gene expression. n=636 DORC gene analyzed. **D**, Schematic diagram of the workflow for identification of transcription regulatory networks [44]. The significance of each TF-gene interaction is tested by assessing whether the TF motif is enriched in the target CRE, and whether the expression of TF correlates with the accessibility of the target CRE (see Methods). **E**, Heatmap of regulation scores for the top 0.1% of TF-DORC enrichments (n=25 TFs and n=147 DORCs). DORC genes are in rows and TFs are in columns. DORCs with CREs overlapping CAD-associated variants are marked with red. Positive and negative regulatory effects are shown in red and blue, respectively. Red arrows highlight the opposite regulatory effect of JDP2 on *CD109* and *PLPP3*. **F**, Network visualization of significant TF-DORC pairs for CAD-DORCs (absolute regulation score > 1.5). Nodes represent CAD-DORCs (orange) and TF regulators (grey). Non-CAD-DORCs regulated by MITF are plotted separately (green discs). Red lines indicate activating effects and blue lines indicate repressive effects. Line widths are weighted by the signed regulation score.

To validate the predicted peak-gene interactions, we tested our macrophage CRE sets for enrichment against a wide range of functional annotations generated from public datasets. By incorporating blood eQTL data from GTEx [68] and eQTLGen [69], we showed that eQTL SNPs (eSNPs) were significantly overrepresented in the CREs (p<0.0001, FigS4A), with at least one eSNP in 41.2% of CREs. The CREs were also enriched for activity-by-contact (ABC) super enhancers identified from enhancer-promoter contacts in mononuclear phagocytes [70] (FigS4B) and for genomic regions shown to interact with macrophage promoters by promoter capture Hi-C [71](FigS4C).

In addition, we quantified the active enhancers signals over all open chromatin peaks using an H3K27ac ChIP-seq dataset we generated in bulk macrophages [39]. The average H3K27ac signal in CRE peaks was significantly higher than in non-CRE peaks (p < 2.2×10^−16^) and exposure to ox-LDL increased the H3K27ac signals in CRE peaks (p = 6.458×10^−6^) but not in non-CRE peaks (Fig4B, FigS4D).

We next applied the FigR framework [44] to the multi-omics data to recover the gene regulation dynamics in macrophages. To further reduce the background noise, we limited the downstream correlation analysis to a subset of 636 genes with a high density of CREs (9 or more CREs per gene, FigS4E). These genes and their associated CREs constitute domains of regulatory chromatin (DORCs). Of these DORC genes, 529 (83.2%) were regulated by strong mononuclear cell ABC enhancers (ABC score>0.1; FigS4F)[70]. Upon ox-LDL exposure, changes in chromatin accessibility at DORC CREs closely correlated with changes in expression of DORC genes (R=0.62, p<0.001) (Fig4C).

For each DORC gene, we assessed their interactions with all detected TFs by two metrics: TF motif enrichment in CRE peaks, and the correlation between TF expression and CRE chromatin accessibility (Fig4D). In total, we analyzed 588,300 TF-DORC pairs representing 925 TFs and 636 DORCs and identified 1612 strong interactions (|regulatory score| > 1.5) involving 95 TFs and 526 DORCs (TableS4). The top 0.1% of TF regulators and their target DORCs with the maximum regulatory scores higher than 2 are plotted in Fig4F. Notably, several TFs, primarily in the AP-1 family, display opposite regulatory effects across different DORC genes (Fig4E). For example, JDP2 has an activating effect on *CD109*, but negatively regulates *PLPP3* (marked with arrows in Fig4E).

Different DORCs that share similar TF associations represent integrated gene regulatory networks. We collated all strong TF-DORC interactions (|regulatory score| > 1.5) with CAD DORCs (DORC with CREs overlapping CAD-associated variants) and constructed a transcription factor regulatory network for CAD in macrophages (Fig4G). This network centered around members of the AP-1 family, with *FOS* and *JUNB* acting as repressors and *FOSL1*, *ATF4*, and *JUND* as activators. Other TF families, including CEBP and KLF, also targeted multiple CAD DORCs. The TF-DORC relationships can be complex; for example, one of the CAD DORC genes regulated by AP-1 factors includes *MITF*, which is itself a TF regulator and activates a wide range of downstream DORCs (Fig4F).

### Prioritization of causal CAD variants

Non-coding disease-associated genetic variants often influence disease risk by altering the gene regulatory properties of CRE [72]. The integration of CRE into the analysis of CAD GWAS results allows candidate causal variants to be linked to their target genes. Permutation-based testing demonstrated that macrophage CREs were enriched for CAD-associated variants—these CREs overlapped significantly more CAD-associated variants than comparable background control peaks did (p=0.0045; FigS5A). The subset of CREs associated with ox-LDL-regulated genes (ox-LDL-response CREs) were also enriched for CAD-associated variants (p=0.0052; Fig5A). We extended GWAS significant SNPs (adjusted p<5×10^−8^) to full summary statistics [18] using stratified linkage disequilibrium score regression (s-LDSC) [50] and showed that this enrichment was specific for CAD compared to non-relevant traits (FigS5B). These results strongly support the hypothesis that macrophages and their response to ox-LDL are causally involved in the heritability of CAD.

**Fig5.**
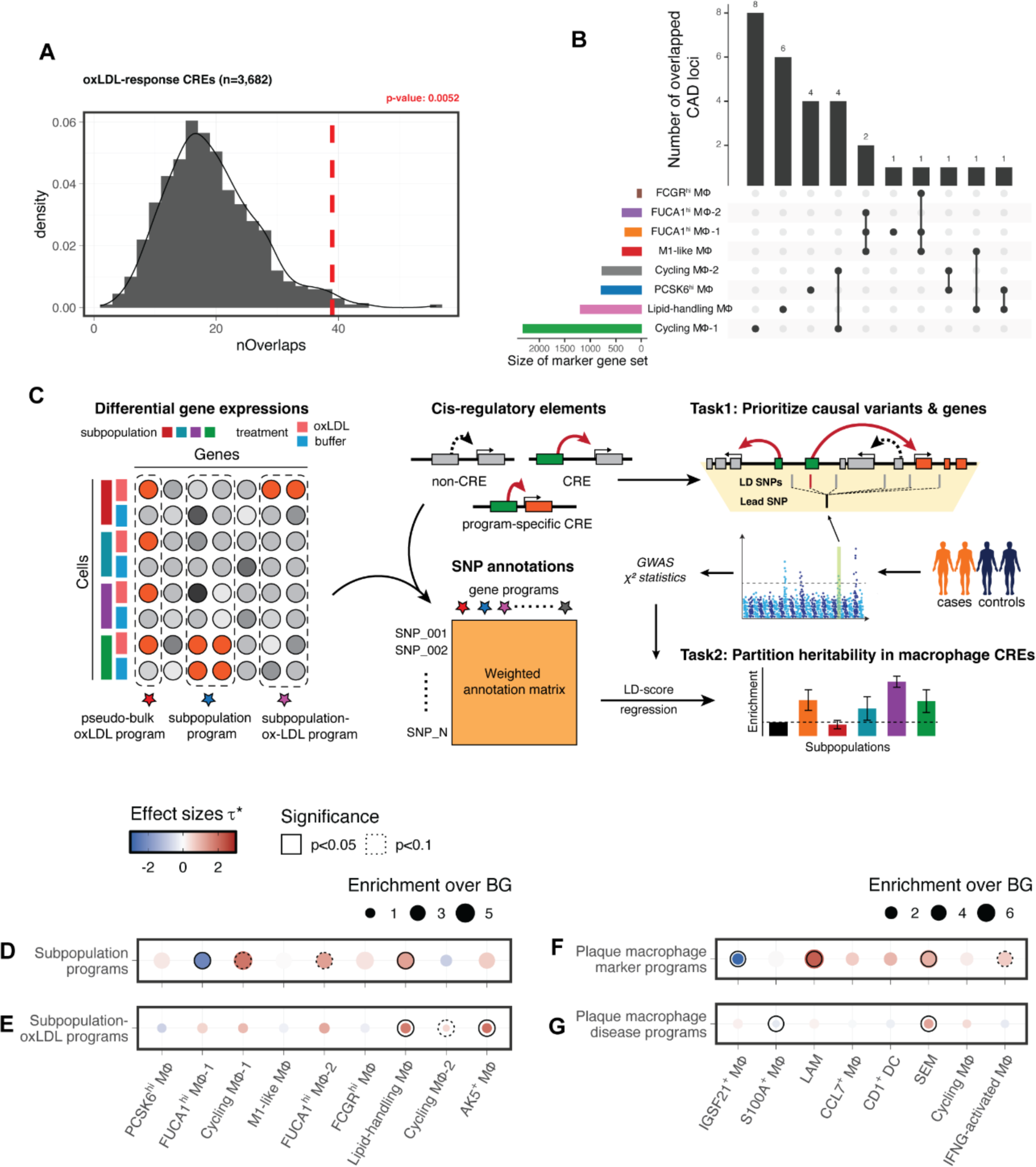
Partitioning the genetic risk of CAD prioritizes disease-critical macrophage subpopulations both *ex vivo* and in plaque. **A**, Histogram showing the null distribution of overlaps between CAD-associated SNPs and n=1000 sets of background peaks matched for GC% bias and average accessibility for all ox-LDL-response CREs. The observed overlap is marked by the red dashed line. **B**, Number of independent CAD loci mapped onto CREs targeting subpopulation-specific genes. Each set of subpopulation marker genes was binarily selected by a differential gene expression test using an FDR threshold of 0.1. Rows represent marker gene sets and intersections between overlaps across different sets are shown as connected dots. The size of each marker gene set is plotted to the left of the matrix, and the size of each intersection is plotted on the top. **C**, GWAS heritability enrichment framework. Left: Deriving macrophage subpopulation gene programs, pseudo-bulk ox-LDL gene program, and subpopulation-ox-LDL gene programs from single-cell transcriptomic data. Middle: Converting gene programs into weighted genomic regions using the macrophage CRE map (top), then constructing SNP annotation matrices by assigning the corresponding weights to 1000G SNPs overlapping each region (bottom). Right: after accounting for the LD structure within GWAS risk loci, the putative causal variants and their target genes are prioritized by their overlaps with macrophage CREs (task1); the relative enrichment of GWAS heritability for stratified subpopulation-specific / subpopulation-specific-ox-LDL-specific annotations are tested using the LD score regression model [50] (task2). **D** and **E**, Enrichments for CAD heritability in subpopulation gene programs (**D**) and subpopulation-ox-LDL gene programs (**E**). Dot size denotes the relative enrichment over background annotations. Color denotes the standardized effect sizes (see text) after conditioning on each other (**D**) or on a pseudo-bulk ox-LDL-response program (**E**) Nonzero τ* estimates reported at two significance levels (solid: p<0.05, dotted: p<0.1). **F** and **G**, Enrichments for CAD heritability in marker programs (**F**) and disease programs (**G**) in plaque macrophages. Dot size denotes the relative enrichment over background annotations (note the larger scale compared to previous panels). Color denotes the standardized effect sizes after conditioned on each other (**F**) or on a pseudo-bulk disease vs healthy program (**G**). Nonzero τ* estimates reported at two significance levels (solid: p<0.05, dotted: p<0.1).

Of the 9516 variants in high linkage disequilibrium (r^2^ >0.8 in the 1000G EUR panel) with the lead CAD variants in the NHGRI-EBI Catalog, 121 SNPs were located within macrophage CREs and we linked these to 56 cis-regulated target genes (154 SNP-gene pairs; TableS5). In 101 pairs (71.4%), the SNP and cis-regulated gene were >10 kb apart (median distance 42.9 kb), and the cis-regulated gene was the nearest gene to the SNP only in 48 of SNP-gene pairs (34.3%). These results demonstrate the limitations of traditional approaches which prioritize variants by proximity. Furthermore, although 110 of the SNPs (90.9%) within macrophage CREs were eSNPs [68, 69], only 34.3% (53 out of the 154 SNP-gene pairs) had the same target gene for both the CRE and the eQTL effect, suggesting that eQTL analysis alone will not identify the regulatory effects of many risk variants.

We benchmarked our macrophage CRE map against state-of-the-art ABC-enhancer mapping based on promoter-enhancer contact information from 35 public mononuclear cell datasets [70]. While macrophages are known for their involvement in atherosclerosis [5], we found no significant enrichment for CAD risk variants after annotating macrophage open chromatin regions with the ABC-enhancer map (61,950 peaks, targeting 19,969 genes; FigS5C). We also found no significant enrichment for CAD risk variants in the subset of 9,716 ABC-enhancer peaks targeting ox-LDL-response genes (FigS5D). These results suggest that ABC-mapped mononuclear enhancers are limited in their power to prioritize causal CAD variants from background noise in primary macrophages.

Together, these findings demonstrate the value of using single cell multi-omics to generate a CRE map for the prioritization and interpretation of CAD risk variants. This focused approach provides greatly enhanced power and precision for investigating macrophage-specific CAD variants compared to traditional approaches.

### CAD heritability varies in macrophage subpopulations with enrichment in lipid-handling macrophages

As demonstrated above (Fig2A and Fig2B), macrophage subpopulations differ in their transcriptomic and epigenetic profiles. We therefore reasoned that this would make them differ in the extent to which they are functionally influenced by CAD-risk variants and so the nature of their contribution to CAD heritability. More specifically, we hypothesized that the CAD heritability that is enriched in macrophage CREs (FigS5A-B) could be further stratified to evaluate the genetic contribution of each macrophage subpopulation to the disease. In accordance with this hypothesis, of the 44 loci at which CAD SNPs overlap CREs, 29 of these loci overlap CREs whose target genes are only expressed in one or a few macrophage subpopulations (Fig5B), indicating that their disease risk-modifying influence is likely to be specific to certain subpopulations. Since assayed DNA variants with genome-wide significance only explain a small portion of the genetic variance of CAD [73], we again extended the significant variants to all SNPs included in the GWAS summary statistics. For each macrophage subpopulation, we applied s-LDSC to systematically assess the enrichment for genome-wide CAD heritability, rather than the enrichment for individual risk variants (Fig5C).

In principle, the genetic contribution of macrophages can be divided into subpopulations based on their gene expression patterns, and the gene expression patterns reflect both the baseline and the oxLDL-response signatures in the transcriptome. Using our previously published Pi strategy [49], we determined a series of weighted gene sets (gene programs) by scoring differential gene expression statistics for each subpopulation. A gene program characterizes either the baseline expression of a subpopulation (termed as a subpopulation gene program), or the response of a subpopulation to oxLDL (termed as a subpopulation-oxLDL gene program). As a comparator, we pooled all subpopulations and constructed a pseudo-bulk ox-LDL program characterizing the response of the whole bulk macrophage population to ox-LDL.

The gene programs were then converted into weighted genomic regions using our macrophage CRE map; all overlapping 1000G SNPs were assigned the corresponding weight for each region to construct the SNP annotation matrices required for heritability stratification. Heritability enrichment was defined as the proportion of SNP heritability explained divided by the proportion of SNPs included in each annotation matrix. As subpopulation-specific annotation matrices were weighted on a continuous scale, we also computed the standardized effect sizes τ*, defined as the proportional change in per-SNP heritability with a 1 standard deviation increase in the SNP annotation weight, to assess the strength and statistical significance of heritability enrichment (see Methods) [27, 51].

Compared to the background annotation constructed using all macrophage open chromatin regions, subpopulation gene programs show higher enrichment for CAD heritability (Fig5D and FigS5E). This is expected as regulatory elements have been widely reported to be enriched for disease and complex trait heritability [50]. To test the independent contribution of each subpopulation gene program to the overall enrichment for CAD heritability, we further controlled subpopulation gene programs on each other using a multivariate linear regression model. The results showed that CAD heritability was relatively depleted (τ*=-2.24) in FUCA1^hi^ macrophages, and remained significantly enriched (τ*=1.28) in lipid-handling macrophages (Fig5D). This suggests that the baseline expression patterns in lipid-handling macrophages convey higher genetic risks for CAD with respect to other macrophage subpopulations.

While different macrophage subpopulations had distinct subpopulation gene programs (FigS5F), their subpopulation-ox-LDL gene program were highly correlated, indicating a shared overall ox-LDL-response across macrophage subpopulations (FigS5G). Therefore, instead of conditioning on each other, we used the pseudo-bulk ox-LDL gene program as the reference to compute the standardized effect size τ* for each subpopulation-oxLDL program. For most subpopulations, τ* no longer deviated from zero (Fig5E), meaning that the ox-LDL effect in these subpopulations is largely not subpopulation-specific. However, there was still significant enrichment of heritability in the ox-LDL response in lipid-handling and AK5^+^ macrophages, with τ* of 1.81 and 1.97, respectively (Fig5E).

### CAD heritability is enriched in plaque lipid-associated macrophages

We next used our *ex vivo* macrophage CRE map to analyze the CAD heritability associated with different plaque myeloid cell subpopulations. As with our *ex vivo* macrophages, we constructed subpopulation gene programs characterizing marker genes for each myeloid subpopulation in plaques, and converted them into SNP annotation matrices. The resulting annotation matrices displayed distinctive patterns in cycling macrophages, SEM, and LAM populations (FigS6A), reflecting differences in genetic variant involvement in their subpopulation gene programs.

We identified marked enrichments for CAD heritability in marker gene programs in the LAM and SEM subpopulations, with 0.030% and 0.021% of all analyzed 1000G SNPs accounting for 2.73% and 1.67% of the total heritability, respectively. The estimated per-SNP heritability in LAM and SEM subpopulation gene programs was twice as large as that across macrophage CREs, and 7.0-fold and 5.9-fold higher than that across all open chromatin regions in macrophages (Fig5F, FigS6B). After controlling on other subpopulation gene programs, CAD heritability was still significantly enriched in LAM and SEM programs, with τ* of 2.23 and 1.05, respectively (Fig5F). We also found a negative effect size for IGSF21+ macrophages (τ* = −2.89, p=6.6×10^−3^), suggesting their subpopulation gene program is associated with significantly less per-SNP heritability compared to other plaque subpopulation gene programs.

The inclusion of patient-matched control vascular samples [13] in the meta-analysis (FigS6C-D) enabled us to construct disease-specific gene programs by comparing gene expression changes between plaque and control samples for each plaque myeloid subpopulation (FigS6E). Similar to the analysis of *ex vivo* macrophages, we referenced all subpopulation disease-specific gene programs against a pseudo-bulk disease-specific gene program. This revealed the positive contribution of SEM, and negative contribution of S100A+ macrophages (i.e., CAD heritability is relatively depleted in the disease program in this subpopulation), to the overall CAD heritability enriched in disease-specific gene program (Fig5G, FigS6F).

### CAD risk variants within lipid-handling macrophage-specific CREs

Genome-wide CAD heritability was enriched in both the subpopulation and the subpopulation-ox-LDL gene programs only for lipid-handling macrophages compared to other macrophage subpopulations (Fig5D). Therefore, we reasoned that one or more risk variants would exert their disease risk-modifying influence through this subpopulation. Among the variants conveying CAD heritability, 19 variants (at 6 independent risk loci) with genome-wide significance mapped to marker genes of lipid-handling macrophages (Fig5B). These risk variants were present at CREs targeting the *MITF*, *LPL*, *FDX1/RDX*, *BCAR1*, *TEX2*, and *PLPP3* gene. The mechanism of action of one such variant in the *PLPP3* gene was previously characterized by our group based on studies of bulk macrophage populations [39], providing important orthogonal independent validation of the current approach.

*MITF* is a key transcription factor influencing lysosome function and autophagy [74, 75]. An intronic variant rs12714757 has been associated with CAD risk in both European and East Asian populations (*β*=0.033 and 0.048, respectively) [18, 76]. In our analysis, two closely linked SNPs, rs6772383 and rs11452399 overlapped an intronic CRE of *MITF* (Fig6A). At rs6772383, the risk T allele disrupts a TEAD1-binding site and is associated with decreased *MITF* expression in GTEx whole blood eQTL data (Fig6A-B, FigS7A), indicating that *MITF* expression is protective against atherosclerosis. Exposure of cells to ox-LDL resulted in a moderate increase in enhancer histone signals flanking this CRE (Fig 6A) and highly significant increases in both the chromatin accessibility of this CRE (p=3.3×10^−8^) and *MITF* expression (average log2FC=0.39, p<2.2×10^−16^). Ox-LDL-induced expression of *MITF* was confirmed at the protein level using immunofluorescence staining (Fig6C and FigS7B). LAMs within plaques had higher *MITF* expression than both LAMs in adjacent control tissue (Fig6D, FigS7C) and other plaque macrophages (FigS7D).

**Fig 6.**
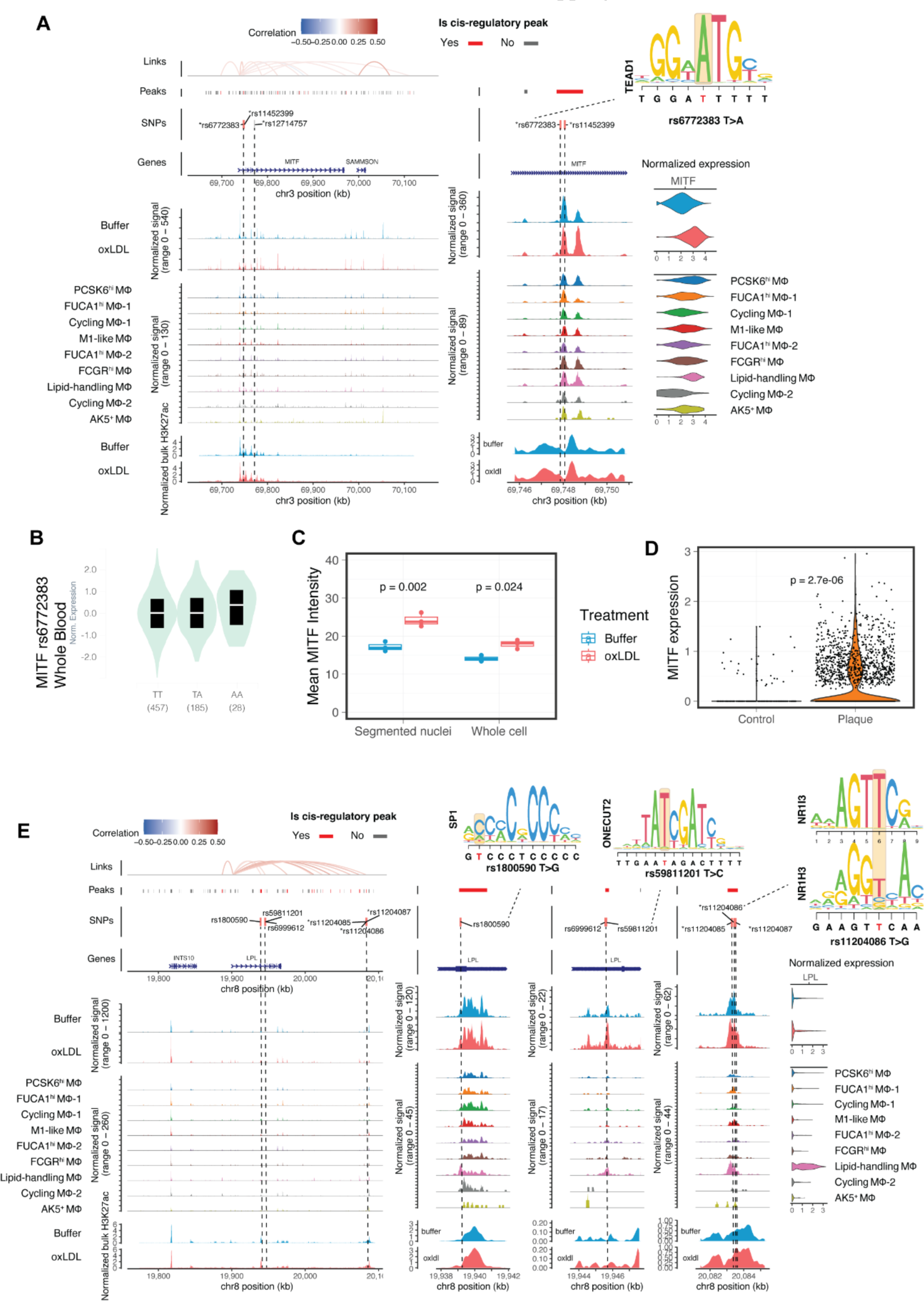
Chromatin regulatory landscapes of CAD risk variants in lipid-handling Macrophages. **A**, Genomic plots for the region encoding CAD risk variants rs6772383 and rs11452399 which localize to a CRE at the *MITF* locus. Tracks (from top to bottom) showing: cis-regulatory interactions (links), ATAC-seq peaks (peaks), CAD-associated variants (SNPs), genomic annotations (Genes), ATAC-seq signals aggregated by treatment, ATAC-seq signals aggregated by subpopulations, enhancer signals from H3K27ac ChIP-seq. Higher resolution coverage plots centered on the cis-regulatory peak are plotted in the middle panel. Violin plots of aggregated gene expression are shown to the right. A red bar indicates significant CREs and eQTL SNPs are marked with an asterisk. **B**, Violin plot showing the allele-specific effect of rs6772383 on *MITF* expression in human blood based on GTEx data. **C**, Quantification of the immunofluorescence staining of *ex vivo* macrophages for MITF, showing oxLDL-induced up-regulation in both nuclear and whole-cell levels (n=3 independent donors). P-values were calculated using t-test. **D**, *MITF* expression in LAMs from plaques vs LAMs from control unaffected arteries. P-value was calculated using Wilcoxon test. **E**, Genomic plot similar to that in (**A**), but showing CAD risk variants rs1800590, rs59811201, rs11204085, rs11204086, and rs11204087 overlapping CREs at the *LPL* locus.

The *LPL* gene encodes lipoprotein lipase which plays a role in lipoprotein uptake and is one of the strongest marker genes of lipid-handling macrophages (Fig3B); its expression is increased by ox-LDL exposure (average log2FC=0.32, p=3×10^−4^). We identified 6 CAD risk variants in 3 different CREs at the *LPL* locus (Fig6E): rs1800590 in a 5’ UTR CRE, rs6999612 and rs59811201 in an intronic CRE, and rs11204085/6/7 clustered in a distal intergenic CRE (*β*=0.091, 0.126 and 0.028 in European population, respectively). Although no significant accessibility change was found in any of the three CREs with ox-LDL exposure (likely because they were already highly accessible), enhancer histone signals increased 70.9% in the CRE containing rs11204086, and 43.0% in CRE containing rs1800590. The protective G allele of rs11204086 disrupts a binding motif for NR1H3 and is associated with higher blood expression of *LPL* in eQTL data (FigS7E). NR1H3 itself is activated notably in lipid-handling macrophages (Fig3G) indicating that the cis-regulatory effect of rs11204086 may be mediated by a subpopulation-specific transcriptional network.

### The CAD-associated variant rs10488763 alters chromatin structure at an AP-1 site and regulation of *FDX1* expression by CEBPB

From the CRE-based variant prioritization analysis, we identified two eSNPs (rs10488763 and rs1443120) in tight linkage disequilibrium (r^2^=1 in EUR panel) that overlap a distal CRE for *FDX1* and *RDX*. This CRE is 56 kb and 77 kb upstream of the transcriptional start sites of the *FDX1* and *RDX* respectively (Fig7A). The alternative alleles of these two SNPs were associated with higher CAD risk (*β*=0.036) [18, 76] and lower expression of both *FDX1* and *RDX* in whole blood (FigS8A). However, CAD summary statistics and eQTL mappings suggest that the GWAS signal aligns most clearly with the eQTL signal of *FDX1* rather than of *RDX* (Fig7B, FigS8B). We also found elevated *FDX1* in plaque LAMs (p<1×10^−4^; FigS8C) compared to non-plaque tissue LAMs, while *RDX* expression remained unchanged (FigS8D). These results indicate that *FDX1* is the likely causal gene influenced by one or both of the CAD variants rs10488763 and rs1443120 at this locus.

**Fig7.**
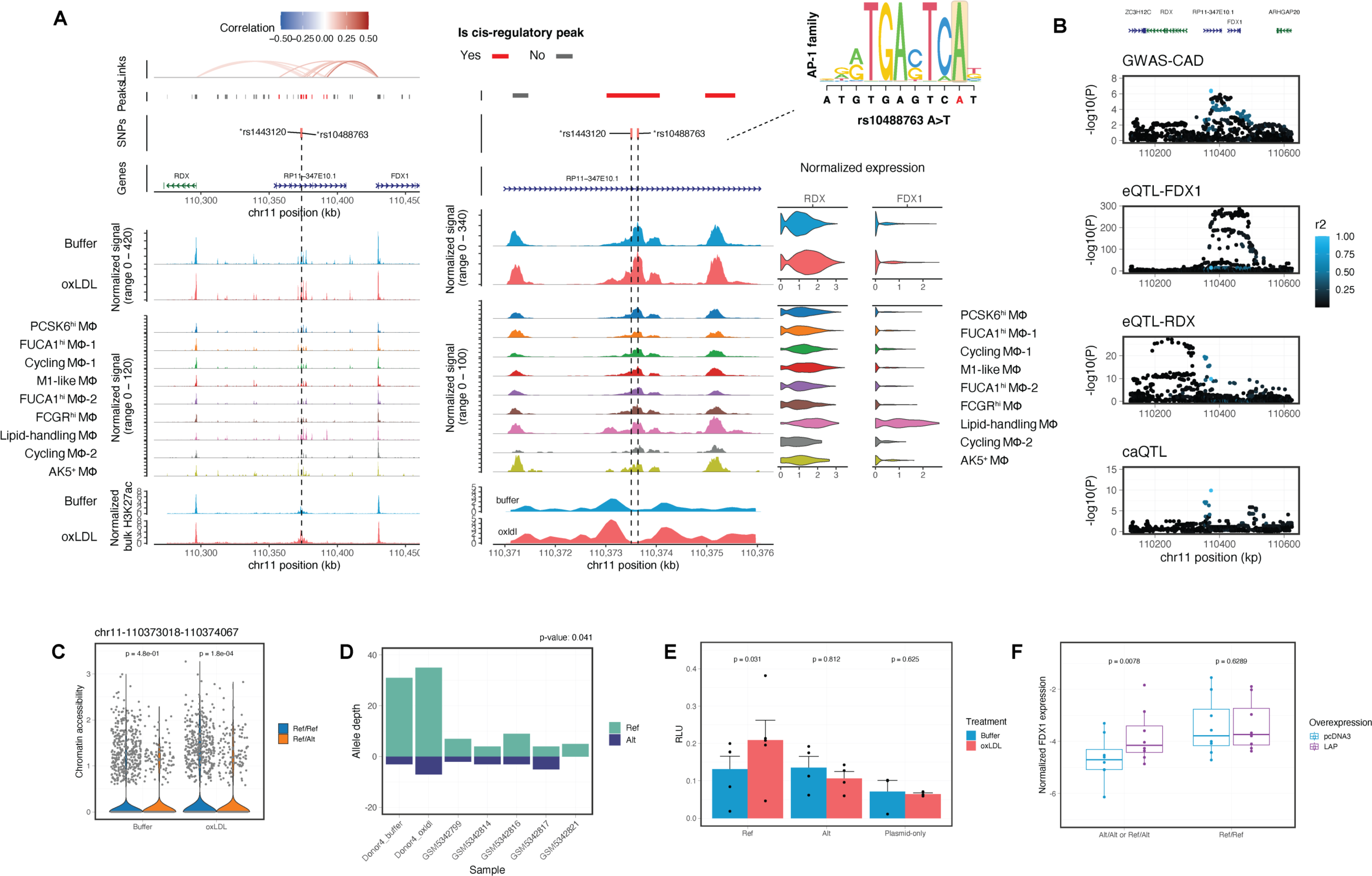
CAD risk variants at the cis-regulatory region of *FDX1* and *RDX*. **A**, Genomic plots of CAD-associated GWAS variant rs10488763 at the FDX1/RDX locus, similar to the plots shown in Fig6A. **B**, Association between rs10488763 variant, CAD (GWAS-CAD), *FDX1* expression (eQTL-FDX1), *RDX* expression (eQTL-RDX, note smaller scale of y axis), and chromatin accessibility (caQTL) at the *FDX1/RDX* locus. SNPs are colored by their LD with rs10488763. **C**, Chromatin accessibility of the cis-regulatory peak in (**A**) grouped by genotype from multi-omics data. (n=3 for A/A, n=1 for A/T). P-values were calculated using Wilcoxon test. **D**, Allelic sequencing coverage of rs10488763 in 24 heterozygous donors, showing higher accessibility in the chromosome containing the protective A allele compared to the T allele. Reads are aggregated from snATAC-seq datasets of primary human macrophages and human plaques [77]. **E**, Luciferase reporter assays in primary human macrophages and ox-LDL-exposed macrophages with using both alleles of rs10488763. (n=5 biological replicates). Signals normalized to Renilla luciferase activities. **F,** RT-qPCR measurement of *FDX1* expression. Primary macrophages (n=8, 6, 2 for Ref/Ref, Ref/Alt, and Alt/Alt samples, respectfully) were transfected with LAP overexpression plasmid or control pcDNA3 plasmid. *GAPDH* was used as the reference gene. P-values shown in **D**, **E** and **F** were calculated using paired Wilcoxon test.

As the risk T allele of rs10488763 perturbs the binding motif for AP-1 factors (Fig7A), we hypothesized that this SNP exerts a cis-regulatory effect by modulating local chromatin structure. The rs10488763 SNP has been linked to a chromatin accessibility QTL locus (caQTL) in iPSC-derived macrophages in an infection model [47] (Fig7B, FigS8B). Consistent with this caQTL signal, we found higher chromatin accessibility in ox-LDL-treated macrophages that are homozygous for the protective A allele, compared to those that are heterozygous for this allele (p=1.8×10^−4^; Fig7C). We further compared the snATAC-seq signal of rs10488763 in heterozygous donors and found that on the chromosome containing the protective A allele, this site was more accessible in both *ex vivo* macrophages and in human plaque cells [77] (p=0.041; Fig7D).

In addition to the chromatin modulating effect of rs10488763, chromatin accessibility at the locus of this variant is also increased by ox-LDL (p=0.015; Fig7A) and this is associated with a 45.3% increase in the local H3K27ac enhancer signal (Fig7A, lower panel). To test if rs10488763 alleles differentially affect this ox-LDL-induced enhancer activation, we transfected primary macrophages with luciferase reporter plasmids containing each allele. We observed a significant allelic difference, with ox-LDL triggering increased enhancer activity with the protective A allele (average 1.58-fold, p=0.031), but not at all with the risk T allele (Fig7E).

Inspection of our CEBPB ChIP-seq data from bulk macrophages [39] revealed a CEBPB binding site 300bp downstream of the rs10488763 SNP within the same CRE (FigS8E). Overexpression of the activator isoform of CEBPB (LAP) induced a 7-fold increase in the enhancer activity of this CRE (FigS8E and F). To test whether the cis-regulatory effect of rs10488763 is mediated by the recruitment of CEBPB, we overexpressed LAP in primary macrophages and quantified the expression of *FDX1* using RT-qPCR (Fig7F and FigS8G). Consistent with the eQTL signals in whole blood samples (Fig7B and FigS8E), *FDX1* expression was significantly lower in *ex vivo* macrophages heterozygous or homozygous for the risk T allele (p=0.0039, Fig7F). Overexpression of CEBPB compensated for this reduction of *FDX1* expression in macrophages carrying the T allele (p=0.016, Fig7F), while having no additional effect in macrophages homozygous for the A allele (p=0.63, Fig7F).

Together, our analyses link the GWAS signal at the rs10488763 locus to *FDX1* and demonstrate an allelic effect on the ox-LDL-induced enhancer activation and change in chromatin accessibility. Our data are consistent with a model in which the interaction of AP-1 transcription factors is altered by variation at this SNP and this influences CEBPB recruitment to the CRE and so upregulation of FDX1 (FigS9). Given that *FDX1* and *CEBPB* were specifically expressed in lipid-handling macrophages (Fig2E-F, Fig7A), our results highlight the crucial role of lipid-handling macrophages as the causal subpopulation with active transcriptional regulatory network required for the CAD risk variant rs10488763 to take effect.

## Discussion

### Beyond current single-cell studies on plaque cells

Large-scale genome-wide association studies have identified many genomic loci associated with CAD, but the molecular mechanisms accounting for these associations are largely unknown. Recent studies using single-cell sequencing approaches on atherosclerotic lesions have provided insights into how these risk loci affect specific cell types like endothelial and smooth muscle cells, yet failed to capture the genetic contribution of macrophages [11, 30, 78, 79]. The numbers of cells available from lesions is typically limited and most are sampled from advanced lesions. In these lesions, the functional pathogenic activity of environmentally derived stimuli, such as atherogenic lipids, may be historic, making genotype-environment interactions difficult to evaluate. For macrophages, these limitations are especially problematic as macrophages are highly plastic and gene regulatory programs are strongly influenced by prevailing stimuli and neighboring cellular activity [47, 80]. To overcome this, we have adapted a simplified model using human macrophages derived *ex vivo* from circulating monocytes. This *ex vivo* model allows us to characterize the transcriptional regulatory dynamics before and after the exposure of the atherogenic lipid, ox-LDL, and comprehensively evaluate the contributions of different macrophage subpopulations to CAD heritability.

### Identification of lipid-handling macrophages

We demonstrate that the transcriptional diversity found among monocyte-derived macrophages mirrors that of the lipid-associated macrophage found in human atherosclerotic plaques. This particular macrophage subset, identified by their active lipid metabolism, is also found in liver and adipose tissue [37, 38] and typically located near blood vessels, the entry points for monocytes into the tissue [58].

Among these monocyte-derived macrophages, we discovered a distinct subpopulation that expresses high levels of cholesterol-clearing genes and exhibit subpopulation-specific transcriptional networks driven by CEBP and NR1H3 family factors. These lipid-handling macrophages can be characterized using the cell surface molecule CD52, and notably accumulate less ox-LDL than other macrophage subpopulations. While direct testing of lipid uptake in human plaque macrophages presents a challenge, the remarkable similarity in transcriptomic profiles between the lipid-handling macrophages and the LPL+ LAMs suggests a potentially significant role for LPL+ LAMs in *in vivo* lipid clearance. These findings may cast light on the rare adverse cardiovascular events including myocardial infarction that have been reported with anti-CD52 therapy (alemtuzumab) [81]. Binding of alemtuzumab to plaque LAMs within arterial walls may trigger intramural inflammation from acute damage to these cells and depletion of CD52+ lipid-associated macrophages from the vessels wall could increase CAD risk.

### AP-1 transcription factors regulate ox-LDL-induced gene expression changes

Our results demonstrate a central role for AP-1 family members in the transcriptional regulation of CAD genes in macrophages, which is consistent with the recent observation that accessible AP-1 motifs were enriched in plaque macrophages [30]. Binding of certain AP-1 subunits, such as JDP2 and FOSL2, was associated with either upregulation or downregulation of gene expression for different genes. AP-1 transcription factors are typically heterodimers of different AP-1 subunits and a given subunit may mediate different gene regulatory function when dimerized with different subunit partners. In addition, AP-1 subunits have been shown to display opposite functions even in the same cell type [82].

### FDX1/RDX locus

*FDX1* encodes an important mitochondrial iron-sulfur protein reductase which is an electron donor for many reducing reactions [83] and is itself reduced by the NADPH-dependent ferredoxin reductase (FDR). Key mitochondrial complexes involved in respiration are regulated by lipoylation which is itself regulated by FDX1 and FDX1 deficiency results in loss of respiration [84]. A genome-wide CRISPR screen demonstrated that it was essential for oxidative phosphorylation [85]. While mitochondrial dysfunction is recognized as a contributory mechanism in the initiation of atherosclerosis [86], the role of ferredoxin has not been characterized in the context of coronary artery disease.

Our computational and experimental results show that the CAD risk T allele at rs10488763 in a distal enhancer region of the *FDX1* gene, increases CAD risk by lowering *FDX1* expression in macrophages. The mechanism of this effect is complex and involves many transcription regulators. Specifically, we demonstrate the role of both AP-1 family factors and CEBPB in modulating the cis-regulatory effect of this CAD SNP. While the AP-1 binding motif is required for ox-LDL-induced enhancer activation, the expression of *FOS* negatively correlates with the expression of *FDX1*, as suggested in our regulatory network analysis (Fig4G). Overexpression of CEBPB bypasses the allelic effect of rs10488763 on enhancer activity and restores *FDX1* expression in macrophages with the disrupted AP-1 binding site. These results indicate a model in which AP-1 binding at the locus does not directly affect *FDX1* expression, but rather remodels the local chromatin to create space for other TFs like CEBPB (FigS9). This model is consistent with the observation that AP-1 factors bind collaboratively with cell-type-specific TFs to select enhancers and establish accessible chromatin [87].

### CAD variant prioritization in macrophages

The analysis of CREs offers a powerful route to further understand CAD heritability. The macrophage CREs that we identified account for 11.5% of the total CAD heritability and contain 121 risk variants with genome-wide significance. The advantages of the CRE approach over eQTL analysis is evident from the small fraction of these 121 variants that have a cis regulatory link to the gene with which they are associated by eQTL analysis. Indeed, multiple studies have now reported limited overlaps between eQTLs and GWAS trait-associated variants [88–90], which highlights the need for other approaches to identify causal genes [91].

Single-cell sequencing approaches circumvent the need for prior identification and segregation of disease-relevant cell types for functional genomic studies [92]. Most single-cell variant prioritization studies integrating single-cell epigenome data only use open chromatin regions, but such regions are not necessarily regulatory [30, 78]. Only 6.9% of open chromatin peaks have cis-regulatory effects in our macrophage data, so colocalization analysis using all open chromatin sites would likely include many false positives for causal variants. A further issue is that the presence of many non-regulatory peaks in accessible chromatin regions drowns out enrichment signals from true regulatory elements, leading to false negative results in prioritization of causal cell types or cell states. This is evident in macrophages for which we have previously profiled open chromatin regions in bulk samples, and found no global enrichment for CAD risk loci [39]. Similar false negatives were also observed in single-cell ATAC-seq studies on human plaque and heart macrophages [30, 78].

In this study, we developed a computational workflow to partition SNP heritability among macrophage subpopulations. Our results demonstrate that CAD heritability is not uniformly distributed across these subpopulations. CAD-associated variants likely act by altering gene regulation, so different macrophage subpopulations with distinct transcriptomes and epigenetic profiles are likely to be differently affected by these causal variants. In this way, the impact of a particular CAD-associated variant may differ between subpopulations. In particular, we show CAD heritability is enriched in the gene program specific to lipid-handling macrophages, and in the gene program specific to the ox-LDL response of this subpopulation. We have provided further orthogonal validation of the relevance of such subpopulation-specific CAD variants at the *MITF*, *LPL* and *FDX1* loci. As cross-validation, we extended this analysis to human plaque macrophages and found that subpopulation-specific gene programs in LAM and SEM cells are also enriched for CAD heritability. To our knowledge, this is the first primary genetic evidence linking these two subpopulations to human coronary artery disease. Our work provides a framework to systematically quantify the disease relevance of heterogeneous macrophage subpopulations and illustrates the substantial value of using single-cell information to improve the genomic resolution of the genetic architecture of human diseases.

## Acknowledgment

The authors would like to thank all healthy volunteers who agreed to participate in this study. We also thank the Oxford Genomic Centre for their support. Illustrations in Fig1B, FigS4D and FigS9 were created with BioRender.com.

